# Progressive chromosome shape changes quantified during cell divisions

**DOI:** 10.1101/2024.11.19.623756

**Authors:** Yasutaka Kakui, Yoshiharu Kusano, Tereza Clarence, Maya Lopez, Toru Hirota, Bhavin S. Khatri, Frank Uhlmann

## Abstract

Mitotic chromosomes give genome portions the required compaction and mechanical stability for faithful inheritance during cell divisions. Here, we record human chromosome dimensions from their appearance in prophase over successive times in a mitotic arrest. Chromosomes first appear long and uniformly thin. Then, individual chromosome arms become discernible, which continuously shorten and thicken - the longer a chromosome arm, the thicker it becomes. The observed chromosome arm length to width relationship can be described by a power law with progressively increasing exponent. In the search for a molecular explanation of this behavior, the popular *loop extrusion* model provides no obvious means by which longer arms become thicker. Instead, we find that simulations of an alternative *loop capture* model recapitulate key features of our observations, including the gradually developing arm length to width relationship. Our analyses portray chromosomes as out-of-equilibrium structures in the process of transitioning towards, but on biologically relevant time scales not typically reaching, steady state.

## Introduction

The process by which mitotic chromosomes arise from apparently amorphous interphase chromatin, in preparation for cell divisions, has captivated cell biologists for a long time^1,2^. Chromosomes bestow genome portions the required compaction and mechanical stability for segregation by mitotic spindle forces^3,4^. While on the one hand constituting stable entities, chromosomes are also known to change their appearance over time. Early observations in plant root cells, in which cell divisions were arrested by the spindle poison colchicine (c-mitosis), revealed how chromosome arms gradually resolve to yield archetypal X-shaped chromosomes^5,6^. Successive chromosome shape changes have also been seen more recently^7–9^. When chromosomes become first discernible in prophase, they appear uniformly thin^10^. At later mitotic stages, chromosome arms have shortened and thickened, with longer arms now wider than shorter arms^11^. Despite the accumulated evidence for chromosome plasticity, a systematic quantitative analysis of chromosome shape changes over time, and an exploration of what these shape changes reveal about chromosome architecture, remains to be performed.

The chromosomal multisubunit protein complex condensin, a member of the Structural Maintenance of Chromosomes (SMC) family, lies at the core of mitotic chromosome formation^12,13^. Condensin introduces a layer of mitosis-specific, long-range chromatin contacts. The span of these condensin-mediated chromatin contacts differs between species. Within small budding yeast chromosomes, they span tens of kilobases. In fission yeast, condensin-dependent mitotic interactions reach hundreds of kilobases, whereas they span megabases in the case of human chromosomes^9,11,14–16^. While the reach of condensin interactions therefore scales with chromosome size amongst organisms, within each species the contact range is invariant amongst chromosomes of different lengths^11^. The molecular mechanism by which condensin establishes mitosis-specific chromatin contacts, and how its species-appropriate contact range is defined, remains incompletely understood^17–21^.

The condensin complex is built around an ABC-ATPase module. ATP binding is required for condensin association with chromosomes, while ATP hydrolysis is necessary to achieve chromosome compaction^22–24^. A popular model posits that the condensin ATPase promotes active extrusion of a chromatin loop, until neighboring condensins meet^25,26^. This ‘*loop extrusion*’ model predicts the formation of a central condensin axis from which chromatin loops emerge. Given similar condensin densities and contact spans along short and long chromosome arms^11^, the model predicts that chromatin loop architecture and therefore chromosome width is the same along short and long arms.

In an alternative ‘*loop capture*’ model for chromosome formation, condensin forms chromatin interactions by sequentially topologically entrapping two DNAs that find each other by Brownian diffusion^27,28^. Simulations of the loop capture mechanism have recapitulated several native-like chromosome features^21,29^. Amongst these features, the model predicts formation of rosette-like chromatin interaction foci, consistent with the experimental observation of punctate condensin clusters in chromosomes^21,30^. Dynamic rearrangement of such chromatin rosettes could allow longer chromosome arms to become wider. However, the predicted dimensions of chromosomes formed by a loop capture mechanism, and how these dimensions might change over time, has not yet been investigated.

Here we quantitatively analyze the shape changes of human chromosomes, from their appearance in prophase over sequential times in a mitotic arrest, paying special attention to the developing length to width relationship. We then compare our measurements to computational loop capture simulations. Initially uniformly thin chromosomes, or simulated chains, continuously shorten and thicken. The length to width relationship of both chromosome arms and simulated chains can be approximated by power laws with an exponent that increases over time. Short chromosome arms, and simulated chains, reach a steady state with a final length to width aspect ratio relatively fast, while longer chains do not reach equilibrium during the times of our observations, or simulations. These considerations depict chromosomes as out-of-equilibrium structures on the way towards, but on biologically relevant time scales often not reaching, loop capture equilibrium.

## Results

### Progressive chromosome shape changes

We synchronized HeLa Kyoto cells at the G2/M boundary by treatment with the cyclin-dependent kinase 1 (CDK1) inhibitor RO-3306. Cells were released from synchronization into medium containing colcemid, allowing mitotic entry but blocking anaphase onset and mitotic exit. Samples were taken at sequential time intervals and processed for chromosome spreading and visualization using the DNA dye 4′,6-diamidino-2-phenylindole (DAPI). Long, thin chromosomes became apparent 12 minutes after G2/M release, which shortened and thickened by 20 minutes (Fig. 1). At 30 minutes, sister chromatids and centromere constrictions became discernible. Sister arms remained cohesed while they continued to shorten and thicken up to the 60-minute mark. By 120 minutes, sister arms separated, remaining connected only at the centromeres, resulting in prototypical x-shaped chromosomes. Over subsequent hourly intervals, chromosome arms continued to shorten and thicken, until we terminated the experiment at 360 minutes. Our observations suggest that, in mitotically arrested cells, chromosomes undergo continuous shape changes.

**Figure 1.**
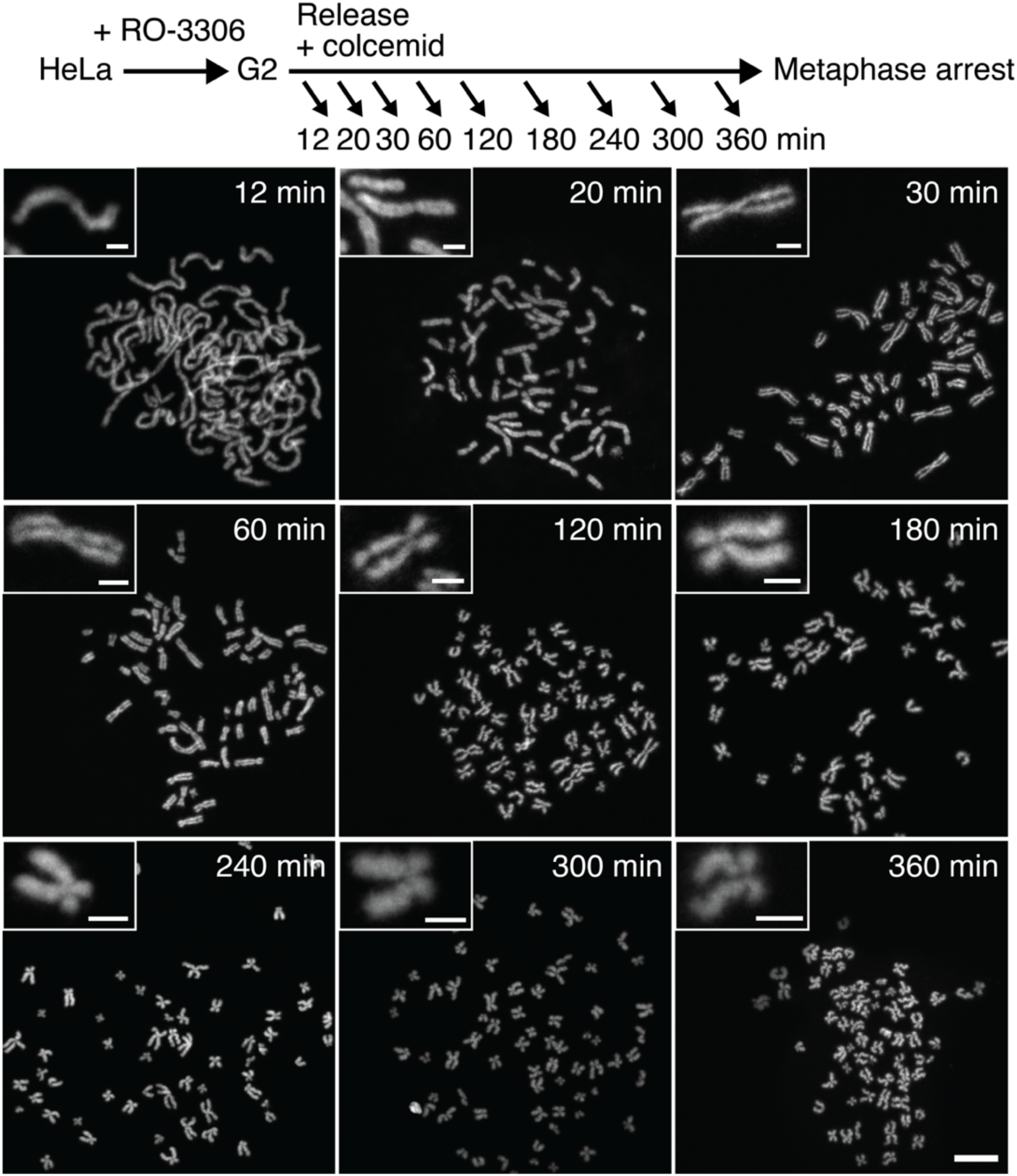
Experimental design and time series of chromosome shape changes. HeLa Kyoto cells were synchronized in G2, released, and maintained in a colcemid-induced mitotic arrest and stained with the DNA dye 4′,6-diamidino-2-phenylindole (DAPI). Example spreads from cells processed at the indicated times are shown. The large images are to scale (scale bar, 10 μm), while the insets show chromosomes at increasing magnifications over time (scale bars, 2 μm).

To determine when chromosome segregation occurs during unperturbed mitotic progression, we repeated the above G2/M synchronization experiment but released cells from the CDK1 inhibitor block into medium without colcemid. We followed mitotic progression using the live cell SiR-DNA stain (Supplementary Fig. 1). For the first approximately 30 minutes, chromosome formation followed a similar trajectory to what we observed in the fixed samples from colcemid-treated cells. Then, unlike in the arrested cells, chromosomes aligned on a metaphase plate, split, and cells exited from mitosis. Anaphase onset occurred at 42 ± 8 minutes (mean ± s.d., n = 50) following CDK1 inhibitor release. Thus, chromosome segregation usually sets in at a time when sister chromatid arms have individualized but remain cohesed.

### Chromosome arm lengths and widths

We applied the following tools to measure chromosome dimensions in the fixed samples from the 360-minute mitotic time course experiment. At the two early time points (12 and 20 minutes), we manually traced the lengths of each chromosome (Fig. 2a). To determine chromosome widths, we selected straight chromosome regions to which we applied a moving Gaussian fit and derived the average full width at half maximum^11^. As centromeres and sister chromatids are not yet discernible, we make the approximation that each of a chromosome’s four arms occupies half the length, and half the width, of a chromosome at these two early time points. We are aware that this approximation underestimates the length of the longest chromosome arms and overestimates the length of the shortest arms.

**Figure 2.**
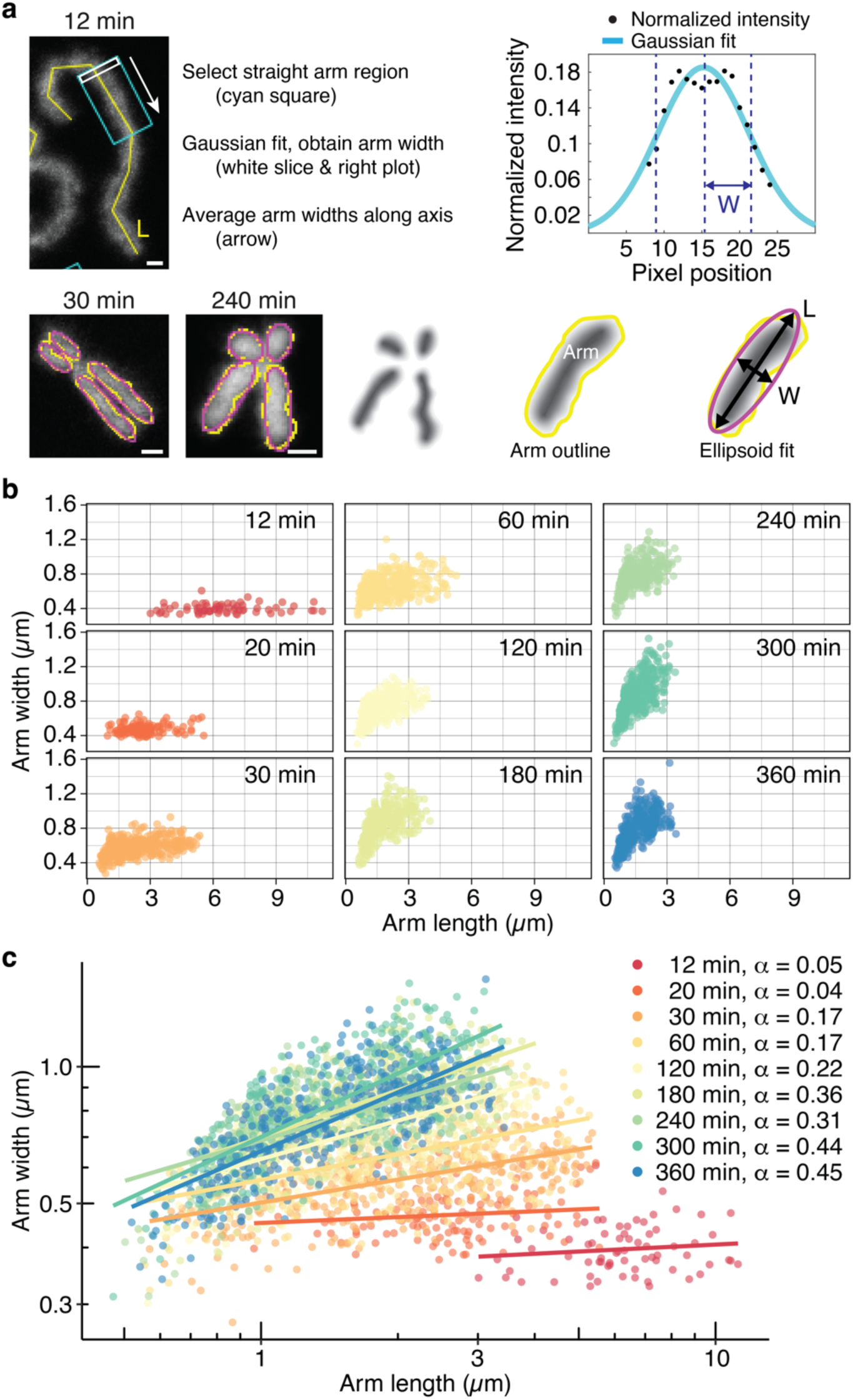
Quantification of the chromosome length to width ratio reveals a power law relationship with an increasing exponent. (**a**) Illustration of the approaches used to measure chromosome arm lengths and widths at early (12 and 20 minutes, top) and later (30 to 360 minutes, bottom) time points (scale bars, 1 μm). (**b**) Chromosome arm widths as a function of arm length. All chromosome arms were measured and are aggregated from two cells at the indicated times. (**c**) Arm lengths and width at all above time points plotted on a double logarithmic scale, with power law fits and power law exponents indicated.

From 30 to 360 minutes, individual chromosome arms are visible, and we traced them using a manual threshold, emulating the half maximum intensity threshold used to determine chromosome width at earlier times. To measure arm dimensions, we computationally fitted ellipsoids with the same area as the traced region, then recorded ellipsoid lengths and widths (Fig. 2a). To comprehensively sample chromosome behavior, we measured all chromosome arms from two cells at each time point and aggregated the measurements. Chromosome arms have varied shapes, especially at the latest 360-minute time point when curly chromosome arms emerged. Ellipsoid fitting is therefore a simplification that will sometimes overestimate width, or underestimate length. Our records are not, therefore, meant as an accurate absolute measure of individual arms. Rather, fitting allows us to sample all chromosome arms and deduce overall trends of chromosome shape changes over time. This approach can be justified, as we made qualitatively similar observations about the chromosome arm length to width relationship when using an accurate chromosome width tracing algorithm, which however restricted the analysis to a subset of chromosome arms (Supplementary Fig. 2)^11^.

Plotting chromosome arm lengths and widths over time revealed progressive chromosome arm shortening and thickening (Fig. 2b). At early times (12 and 20 minutes), width was uniform and independent of length. As soon as individual chromosome arms become discernible (30 minutes), longer chromosome arms were wider, a trend that became more pronounced as time progressed.

We previously described the mitotic chromosome arm length to width relationship at the 30-minute time point using a power law (*w* = c *L*^α^, where *w* is the chromosome arm width, *L* is the length, c is a constant and α the power law exponent)^11^. To examine whether power laws describe the length to width relationship across the time series, we plotted our measurements on a double logarithmic scale (Fig. 2c). A linear distribution of measurements on this scale is indicative of a power law relationship, with the slope of a linear fit reflecting the power law exponent. At early times, when width is invariant between chromosomes, the power law exponent is essentially zero. At successive times, the slope, and thus the power law exponent, increased until reaching 0.45. Below, we will further analyze this developing length to width relationship.

Note that a proper test of a power law relationship requires both quantities, here chromosome arm length and width, to vary over several orders of magnitude. This requirement is unavailable in our experiment, so here we use the power law merely as a convenient mathematical form, which captures the empirically observed behavior.

### Short arms equilibrate faster

We next plotted chromosome arm lengths over time. As our fixation protocol does not allow us to track the progression of individual arms, we instead stratified all measured arms by their length at each time point. We then record the length of every 10^th^ percentile, i.e. the length of the 10^th^, 20^th^, etc., until 90^th^ percent longest arm at each time. This approach enabled us to follow the behavior of these percentile lengths over time (Fig. 3a). The approach revealed that the shortest percentile arms rapidly shortened within the first 30 minutes, and after that maintained a relatively constant length. In comparison, the longest percentile chromosome arms showed a different behavior. Shortening also began at a rapid initial rate until 30 minutes, but then gradually continued until the end of our time course at 360 minutes. Intermediate percentile arm lengths show a behavior intermediate between these extremes. Because of the uncertainty around actual chromosome arm lengths in the first two time points (which in the absence of a centromere constriction we approximated as ½ the chromosome length), we did not quantitatively fit a mathematical description to the observed time-dependent length changes. Nevertheless, it becomes qualitatively apparent that short chromosome arms reach an equilibrium length relatively quickly, and before the time when anaphase normally occurs at around 42 minutes. Longer arms, in contrast, are still on the way to an equilibrium by the time of anaphase onset, and even at the end of our mitotic arrest time course.

**Figure 3.**
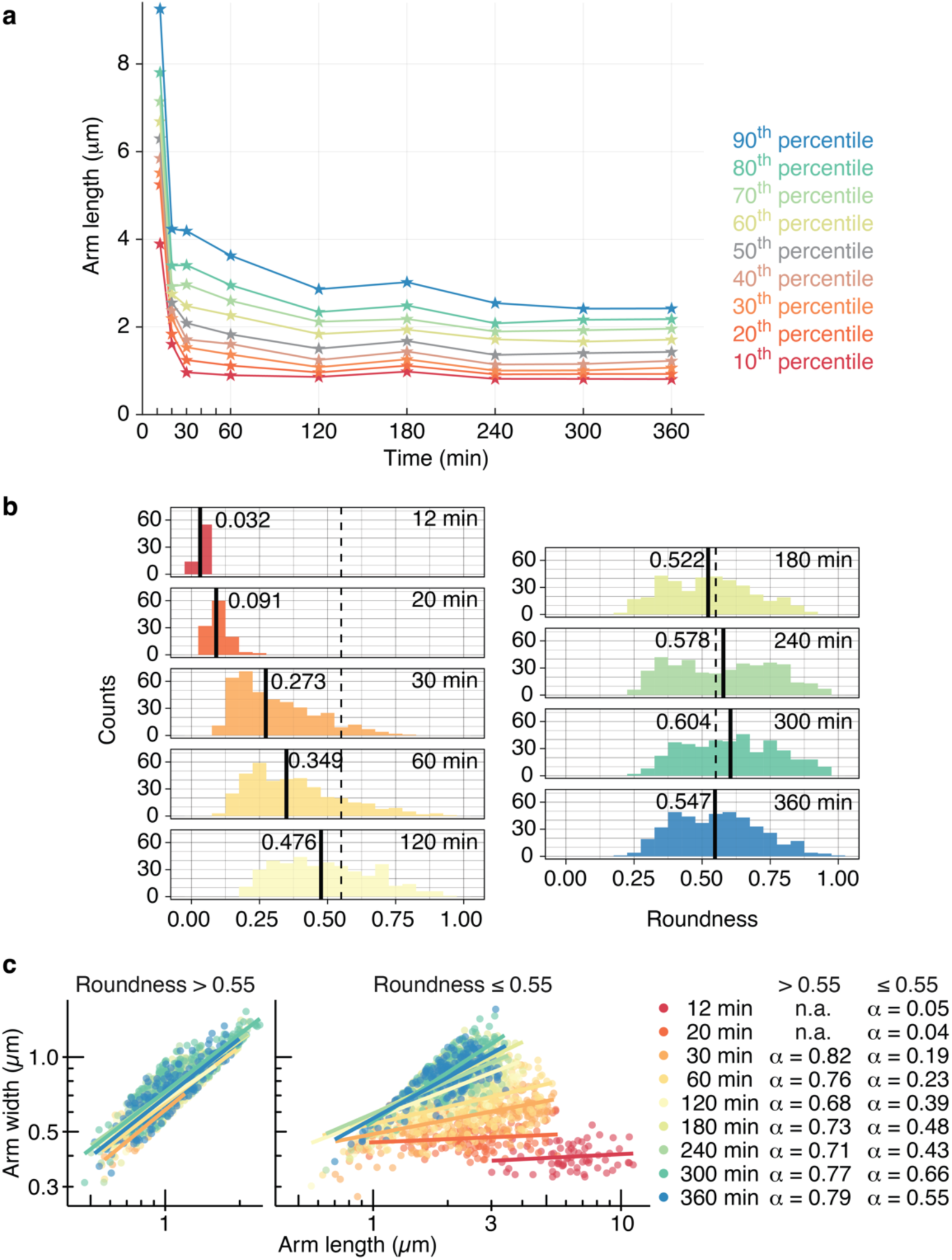
Short chromosome arms reach equilibrium faster than long arms. (**a**) Chromosome arm lengths, divided into 10^th^ percentiles, were plotted over time. (**b**) Chromosome arm roundness (width divided by length) as a function of time. Median roundness at each time is indicated by a solid black line. A roundness of 0.55 is highlighted by a dashed line, corresponding to the mean roundness at the 360 minutes time point. (**c**) Chromosome arm widths as a function of arm length over time, split into arms with roundness greater than or equal or smaller than 0.55. Power laws were fitted to both sets and power law exponents are indicated. n.a., no arms with roundness greater 0.55 were found at the first two time points.

### A final chromosome roundness

Having realized that shorter chromosome arms reach a final equilibrium in our time series, we wanted to characterize the dimensions of this equilibrium state. We therefore looked for a way to distinguish short, equilibrated arms from longer arms. For that purpose, we plotted chromosome arm roundness over time, where roundness *r* is the ratio of arm width divided by arm length. Starting from a very small value of ∼ 0.03 at 12 minutes, roundness gradually increased and spread out over time, reaching an average of ∼ 0.55 at late times (Fig. 3b). Overlaying roundness onto our length to width relationship plots confirms that shorter arms generally reach greater roundness, and sooner (Supplementary Fig. 3). We then used the final median observed roundness at 360 minutes of *r* = 0.55 to divide our chromosome arm distribution. Plotting all chromosome arms with a roundness of greater than 0.55 shows that their length to width relationship now follows a power law with an exponent of ∼0.75, irrespective of the time when these chromosome arms were encountered in our measurements (Fig. 3c). Thus, the final chromosome arm length to width equilibrium can be described by this power law exponent.

On the other hand, chromosome arms with a roundness ≤ 0.55 display a length to width relationship with increasing power law exponents over time (Fig. 3c). This observation confirms that longer chromosome arms continue to change shape until the end of our observation period, with the power law exponent of their length to width relationship approaching, but not reaching that of short, equilibrated chromosome arms.

### Dimensions of a theoretical and of a simulated polymer

To explore a possible underlying mechanism for the observed chromosome arm width progression, we turned to polymer simulations. We previously used a biophysical model of a chromatin chain to explore how loop extrusion or loop capture interactions differentially affect predicted chromosome properties^21^, and we now repurpose this model to study the resultant chromosome dimensions. We use a coarse-grained chromatin chain consisting of beads, each modeled to represent a ∼2 kb region encompassing ∼10 nucleosomes. Beads are connected by springs, with Brownian dynamics determining the stochastic forces on every bead. A soft repulsion term is applied when beads overlap. We first investigated whether this representation of a self-avoiding Rouse polymer chain adopts shapes that display theoretically expected behavior and dimensions.

Fig. 4a shows representative snapshots of the simulated conformations for polymers of increasing chain lengths (increasing bead numbers, *N*), whilst in equilibrium. The conformations are non-compact and grow in size with increasing *N.* Polymer theory predicts that a random polymer with excluded volume grows in size as *N^v^* (*v* = 0.588, this power law relationship links chain length to polymer size. It is different from the previously discussed power law relationship between measured chromosome length and width, for which we use the exponent α)^31^. When we plot the average measured equilibrium length and width of our simulated chromatin chains as a function of chain length *N* (Fig. 4b), we see that both measures follow these theoretical expectations closely.

**Figure 4.**
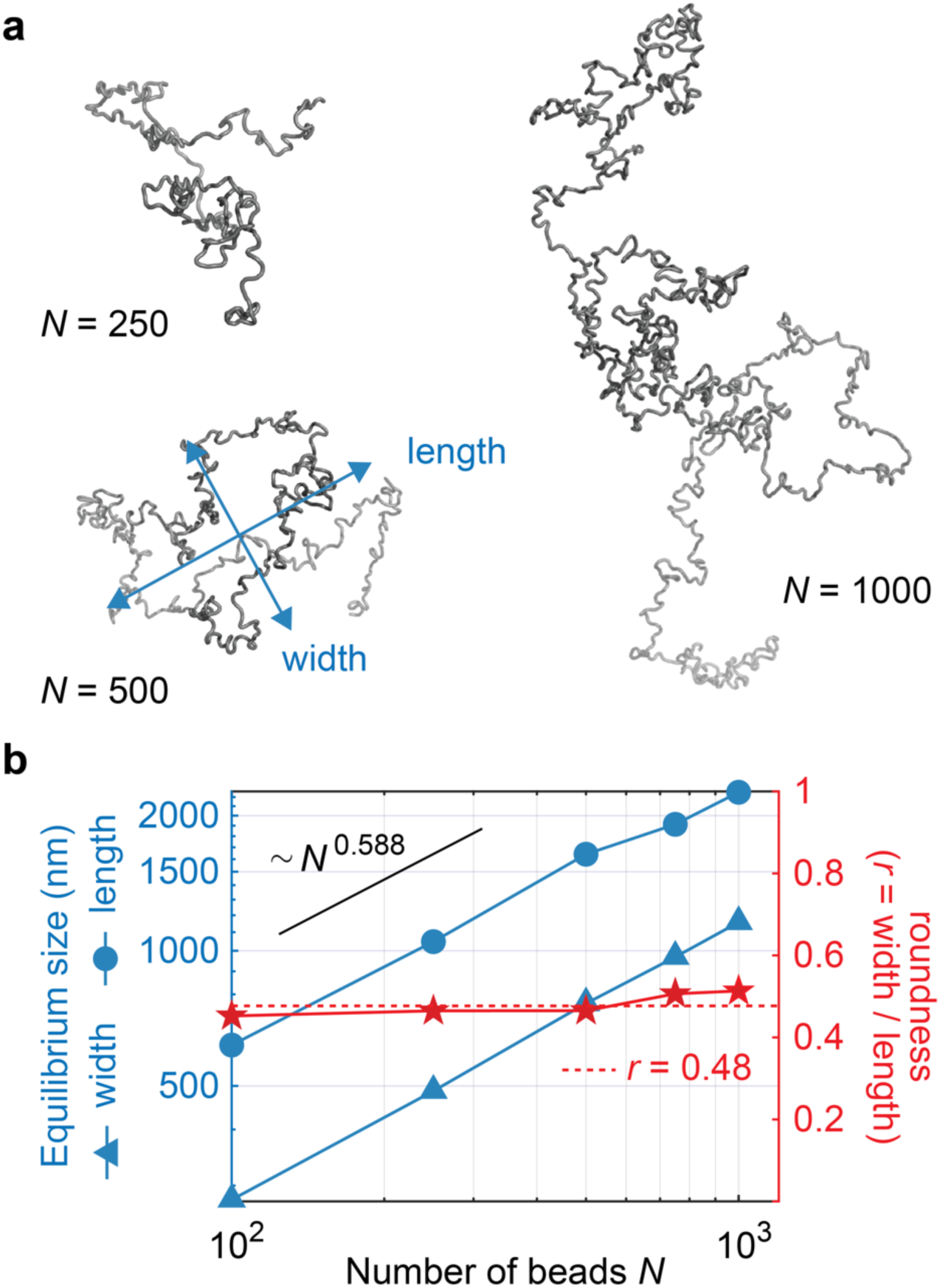
A simulated self-avoiding Rouse polymer, representing an unconstrained chromatin chain, displays a theoretically expected scaling behavior and length to width ratio. (**a**) Snapshots of equilibrated simulated chromatin chains of varying lengths. (**b**) Equilibrium length and width, as well as the width to length ratio, are plotted as a function of chain length. The theoretically expected scaling behavior of random self-avoiding polymers is indicated, as well as their expected roundness (*r* = width / length).

Next, we record the length to width relationship of the simulated chains. Because of the unpredictability of random walks in each of the three available dimensions, the resulting polymer shapes are never perfectly spherical. Based on probability theory, the length, width and depth of a random polymer should be in proportion length : width : depth = 3.44 : 1.64 : 1^32^. If we define the ‘roundness’ of the theoretical polymer as its width divided by its length, in analogy to our experimental chromosome arm measurements, the expected polymer roundness based on these proportions is *r* = 0.48. When we now measure the roundness of our simulated chromatin chains, defined as their equilibrium width divided by their equilibrium length (Fig. 4b), we again see that the simulated roundness matches the theoretical prediction closely.

We conclude that, in the absence of any loop interactions, our simulated chromatin chain adopts a roundness of ∼0.48, as theory predicts. An unconstrained, self-avoiding polymer is therefore on average more elongated than observed chromosome arms in equilibrium, which we have seen above have a roundness of ∼0.55.

### Loop capture interactions shorten the polymer

We next investigated how loop capture interactions impact on the dimension of our simulated chromatin chain. To model loop capture, every 10^th^ beads is selected to be a condensin binding site, corresponding roughly to the empirically observed spacing of ∼23.4 kb/11.7 beads between condensin binding sites in fission yeast^21^. Although condensin binding sites along human chromosomes are spaced much further apart (see below), we expect – as is common in polymer physics – that the scaling behavior from these simulations will be qualitatively representative of the behavior also of much larger structures. Here, we begin by simulating the behavior of fission yeast-sized chromatin chains for computational practicality. If two such condensin binding sites encounter each other by stochastic movements, a pairwise interaction forms and persists with a defined probability, before it turns over. Simulated loop capture results in the emergence of far more compact structures, seen in representative snapshots of equilibrium conformations found for chain lengths *N* = 250 to 2000 (Fig. 5a), when compared to the more extended structures of random walks (Fig. 4a). This compaction is mediated by the formation of characteristic rosettes that constrain the chromatin chain, as also seen in previous loop capture simulations^21,27^. Analyzing the average size of the resultant structures as a function of chain length *N*, we find a similar scaling behavior as for a self-avoiding Rouse polymer (Fig. 4b). Therefore, while more compact, loop capture results in structures that scale similarly to unconstrained random walks with respect to chain length.

**Figure 5.**
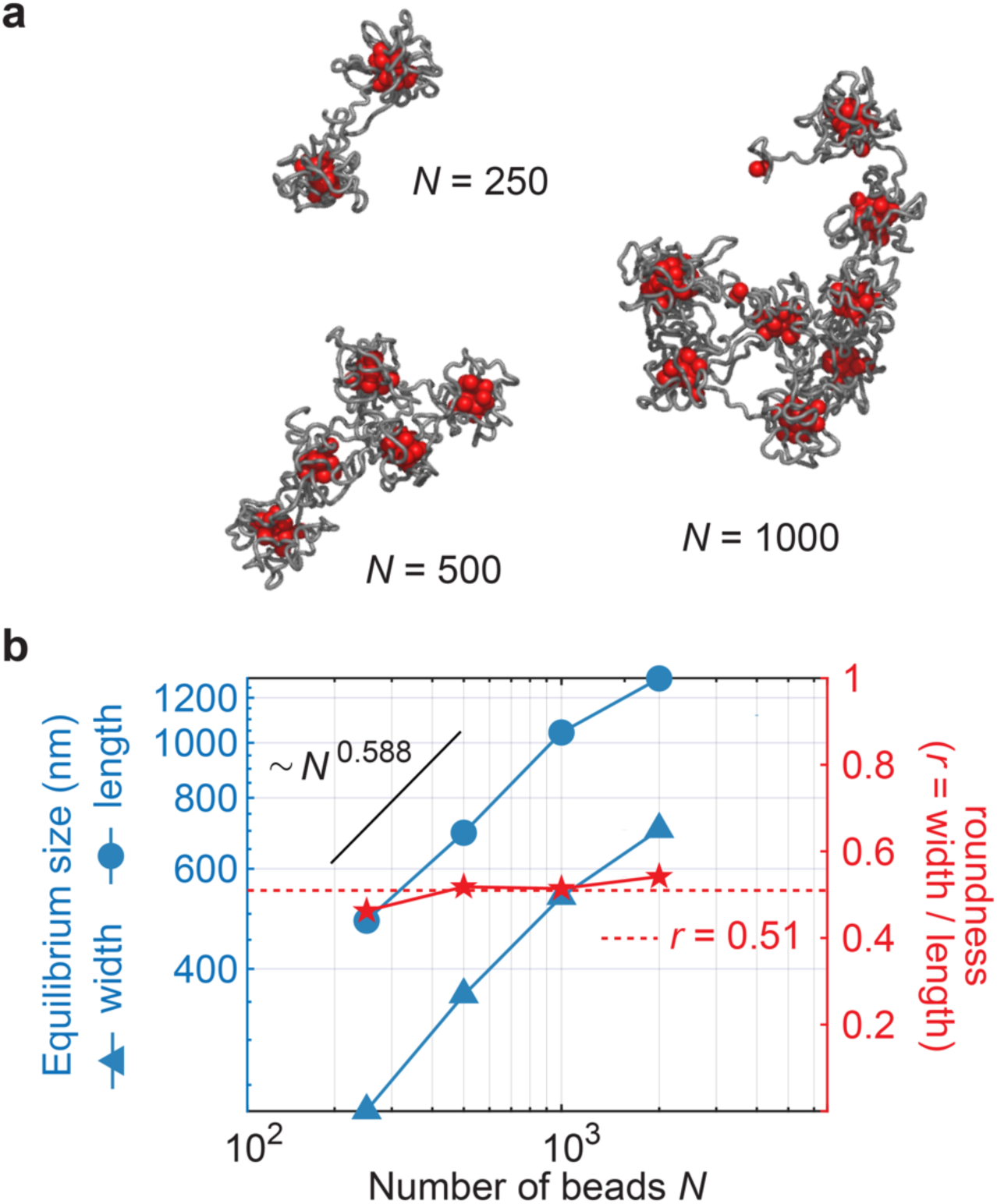
Loop capture interactions compact the polymer and increase its roundness. (**a**) Snapshots of equilibrated simulated chromatin chains on which condensin (red spheres) engages in loop capture interactions. (**b**) Equilibrium length and width, as well as the width to length ratio, are plotted as a function of simulated chain length. The theoretically expected scaling behavior of random self-avoiding polymers is indicated for comparison. The mean observed roundness (*r* = width / length) is highlighted by a dashed red line.

However, when we analyze the width to length ratio of these new, more compact loop capture structures, we find that the mean width to length ratio has increased, corresponding to structures with a greater roundness *r* ≈ 0.51, when compared to an unconstrained self-avoiding Rouse polymer. These findings suggest that loop capture interactions reduce the polymer length more than the polymer width. The new roundness value approaches, but does not completely match, the roundness that we observed for short human chromosome arms that have reached equilibrium during our mitotic time course experiment (*r* ≈ 0.55, Fig. 3b). These observations are consistent with the possibility that loop capture interactions contribute to shaping mitotic chromosomes, but they also suggest that additional mechanisms exist that further increase roundness, at least of short chromosome arms.

The lengths of our simulated chromatin chains are in the Mb range, corresponding in size to fission yeast chromosome arms, and we simulated a condensin binding site interval as observed in fission yeast. In comparison, human chromosomes arms are an order of magnitude longer, in the tens of Mb range. Equally, condensin binding sites are found at approximately ten times greater intervals^21,33^. The scaling behavior of a Rouse polymer with excluded volume, where size grows as *N^v^* (*v* = 0.588), means this behavior is scale free, i.e. the ratio of the size of two polymers only depends on the ratio of their lengths. We observe that following introduction of loop capture interactions this scaling behavior remained essentially unchanged. We therefore postulate, but do not know for certain, that loop capture interactions affect human chromosomes of much larger dimensions in similar ways as seen for our smaller simulated chains.

### Simulated chromosome lengths and widths over time

We have so far considered simulated chromosome arms at their equilibrium. To conclude, we followed simulated chromosome shape changes over time. To do so, we defined an elongated initial state for our simulations, akin to that observed for natural chromosomes that appear around four times longer in prophase when compared to late mitotic stages (Fig. 2b). We initialized chain conformations with a random walk 4x longer than the equilibrium length. Upon release into simulations with loop capture interactions, these elongated chains reached a similar equilibrium state as chains started from an unstretched initial conformation (for chain lengths up to *N* = 2000; Supplementary Fig. 4). Thus, an elongated initial state does not alter the eventual equilibrium chromosome conformation.

We now sampled chromosome conformations over time. Plotting chromosome length as a function of time recapitulated several aspects of the behavior observed of native chromosomes. Short chromatin chains quickly shortened and reached their equilibrium state, while longer chains took longer to reach equilibrium. The longest chains (*N* = 4000) had not reached equilibrium at the end of our simulations, similar to what we observed for long chromosome arms during our mitotic arrest time course. In Fig. 6a, we display representative snapshots over time, for three different chain lengths. These show visually that the shortest chain (*N* = 250) reaches a compact equilibrium structure within 1 minute. For an intermediate size chain (*N* = 1000) it takes roughly 10 minutes to reach a compact equilibrium, while the longest chain length analyzed (*N* = 4000) remains in a visibly non-equilibrated and elongated state at our last, 30 minutes, time point. We quantify this polymer chain behavior by plotting a time series of observed lengths over time for all chain sizes (Fig. 6b, top), which we find is well explained by an exponential relaxation process. From these fits we calculate the relaxation time (Fig. 6b, bottom), which shows an increasing relaxation time with chain length *N*. Qualitatively, the relaxation behavior resembles that found for the relaxation of native chromosomes. Short chromosome arms equilibrate quickly, while longer arms are still in the process of approaching equilibrium at later times.

**Figure 6.**
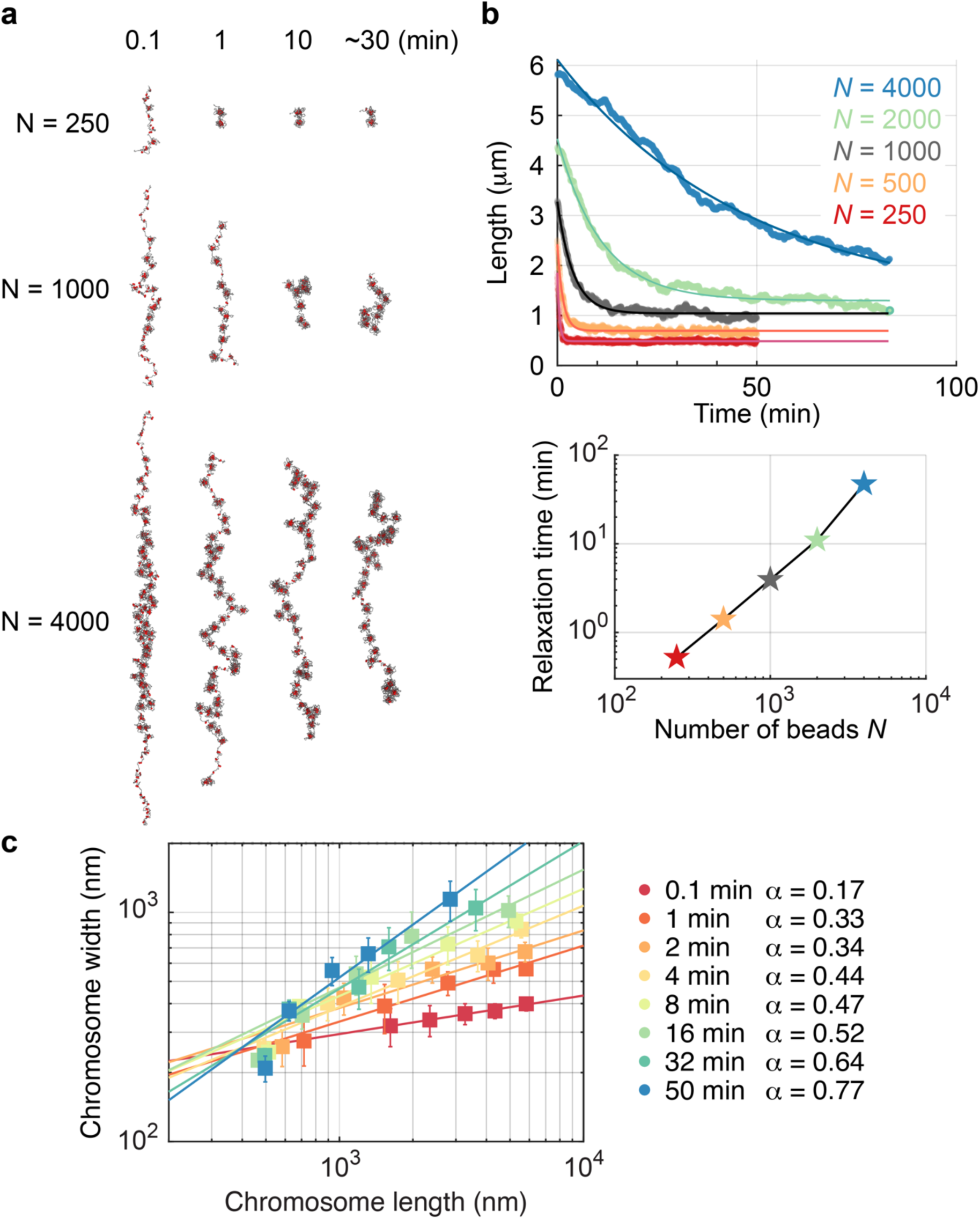
Simulated loop capture interactions result in a gradually increasing power law exponent of the chromosome length to width relationship. (**a**) Snapshots of simulated chromatin chains released from an initially elongated configuration (4 x equilibrium length), on which condensin engages in loop capture interactions. (**b**) Simulated chromosome lengths were plotted over time including exponential fits (top). The exponential fits were used to derive relaxation times, the values of which are plotted alongside as a function of chain length (bottom). (**c**) The chromosome length to width relationship over time. Each data point is the average of 10 replicate simulations at the given time point, with error bars representing the standard error on the mean (errors in length are smaller than the size of symbols used). Power law fits to the length to width relationship are included, and the power law exponents are listed.

Finally, we examine how the simulated chromosome length to width relationship evolves over time, plotting the measured lengths and widths of simulated chains of the five different lengths *N*. Qualitatively, the simulated loop capture chromosomes recapitulate the observed behavior of native chromosomes, where longer chromatin chains become increasingly wider over time (Fig. 6c). When we analyze the measured length to width relationship, similar to what we did for native chromosomes in Fig. 3c, we find that the chromosome length to width relationships can be described by power laws whose exponents increase over time. For simulated chromosomes, the final equilibrium power law exponent approaches ∼0.75, a value that is the same as the exponent that we found describes shorter native chromosome arms that have reached equilibrium during our mitotic arrest time course experiment.

We conclude that a simulated chromatin chain on which loop capture interactions take place, beginning from an elongated state, undergoes shape transitions similar to those observed for native chromosomes. Shorter chains equilibrate faster than longer chains, with developing lengths and widths that can be described by a power law relationship with increasing exponent. These considerations put forward loop capture as a plausible mechanism that contributes to the continuous chromosome shape changes that can be observed of native chromosomes in mitotically arrested cells.

## Discussion

Our study portrays chromosomes in a fresh light. Instead of stable entities, we describe chromosomes as out-of-equilibrium structures, only the shortest of which will have reached their final shape by the time chromosomes split at anaphase onset. We also find that a *loop capture* mechanism provides a plausible explanation for the shape-shifting chromosome behavior, and for a length to width power law relationship whose exponent increases over time, meaning that longer chromosome arms turn successively wider.

Chromatin loop formation by loop capture results in rosette structures, and these rosettes gradually remodel to reach a steady state equilibrium that is defined by the principles of polymer physics. Longer chromatin chains occupy a volume that is not only longer, but also wider. Rosette remodeling appears to be a relatively slow process, unsurprising given the large dimensions of these objects. The process will be influenced by the half-life of condensin-dependent loop interactions that underpin these structures^24,34,35^. It is likely that, additional to rearrangement of existing rosettes, other cellular parameters change over time while cells are blocked in an extended mitotic state. An increasing condensin concentration, histone modifications, divalent cation concentration changes, as well as the depletion attraction force^36–39^ are all factors that could affect chromosome dimensions. Direct condensin-condensin interactions also contribute to vertebrate chromosome formation^40,41^, which were omitted from our simulations. Taken together, a multitude of additional forces impact chromosome formation, and they are a possible reason for why the final observed roundness of human chromosome arms is greater than what we observe in our simple loop capture simulations. Future studies will be required to integrate these additional forces, alongside loop capture, when aiming to fully recapitulate observed chromosome behavior.

A characteristic of chromosome arms that consist of self-organizing chromatin rosettes is their gradual rounding. Rather than forming strictly cylindrical objects, rosettes arrange themselves in an elliptical volume where the middle is wider than both ends. Our simple definition of roundness, as the ratio of width divided by length, did not differentiate between cylinders and ellipses. Based on cursory visual inspection, at least shorter chromosome arms that have reached equilibrium are well described by ellipsoids. An additional hint that chromosome arms show ellipsoid rather than cylindrical architecture comes from chromatin interaction patterns recorded in Hi-C experiments. Chromatin contacts span greater distances in the middle of fission yeast chromosomes arms, when compared to interactions recorded towards both ends^16^, as expected from an ellipsoid. It will be interesting to explore Hi-C datasets from metazoan mitotic chromosomes for signs of an ellipsoid shape.

While dynamic loop capture is a plausible mechanism to explain continuous shape changes, starting from an elongated appearance, an open question remains as to the origin of the initially extended chromosome state. Prophase chromosomes at first are much longer than the outlines of interphase chromosome territories from which they derive^42^, or than the expected shape of an unconstrained polymer chain. The origin of the elongated initial state therefore remains to be understood. Loop extrusion by condensin could generate an elongated bottle brush-like structure with a central condensin backbone^25^, however the widely scattered condensin distribution that is seen in prophase chromosomes does not lend support to this scenario^30^. Short-range chromatin interactions that are established by the cohesin complex might alternatively contribute to generating an elongated shape, or the reptation behavior of neighboring chromosomes within nuclear confines.

In higher eukaryotes, two condensin complexes, condensin I and condensin II, together shape chromosomes^43,44^. Condensin I majorly affects chromosome width, while condensin II predominantly affects chromosome length. How two condensin subtypes exert apparently selective compaction in two orthogonal directions remains unknown. What is known is that condensin II engages in much farther-reaching chromatin contacts while condensin I adds shorter-range interactions^9^. In a loop capture scenario, we can envision how condensin II sets up a coarse rosette architecture, with condensin I inserting a layer of finer-grained rosettes. The larger condensin II rosettes will define the overall chromosome outline, while condensin I compacts internal as well as surface chromatin loops. In this scenario, condensin II is the main driver of compaction, explaining why chromosomes appear more prophase-like, or longer, in its absence. By compacting surface loops, in turn, condensin I might affect chromosome width more than length. Simulating chromatin behavior that arises from two such distinct types of loop capture interactions provides fertile ground for future investigations.

We close by noting that, while displaying vast size differences, chromosomes from across flowering plants and vertebrates all follow a universal length to width relationship^45^, suggesting that they are governed by a common physical principle. Much remains to be learned about the physical and molecular concepts that form these beautiful structures.

## Methods

### Cell culture, synchronization, and chromosome spreads

HeLa Kyoto cells were cultured in DMEM supplemented with 10 % fetal calf serum, 0.2 mM L-glutamine, 100 U/mL penicillin and 100 μg/mL streptomycin at 37 ℃ in a 5% CO_2_ environment. To study the changes in chromosome morphology, cells were synchronized at the G2/M boundary by the treatment with 9 μM RO-3306 for 3 hours, then washed once and released into medium containing 100 ng/mL colcemid. After collecting cells by mitotic shake off at the indicated times, cells were incubated with a hypotonic buffer (PBS:H_2_O = 1:9) for 5 minutes, and fixed with fresh Carnoy’s solution (70 % methanol, 30 % acetic acid). Fixed cells were dropped onto glass slides and dried. Spread chromosomes were stained with 10 μg/mL DAPI in PBS for 5 minutes and mounted using ProLong Gold (ThermoFisher) antifade reagent. Images were captured using a Zeiss LSM880 microscope using an 63x objective and Airyscan detection. To image undisturbed mitotic progression, following RO-3306 release, colcemid was omitted from the release medium. DNA was visualized using the SiR DNA live stain (Cytoskeleton) and images acquired at 2.5 minute intervals using a CellVoyager CQ1 High-Content Analysis System and a 60x objective. The time from the first appearance of chromosome structure in prophase until chromosome splitting at anaphase onset was counted in 50 cells.

### Measurements of mitotic chromosome dimensions

At the 12- and 20-minute time points, chromosome lengths were measured by manually drawing a line along the chromosome, and arm widths were semiautomatically determined after selecting straight chromosome regions using a modified MATLAB script^11^. First, a mask was created by binarizing signal intensities of the original image. In this step, intensities outside the mask were set to zero. Second, a slice perpendicular to the chromosome length axis was taken from the masked image and a single Gaussian fit was applied to the slice and the full width at half maximum (FWHM) calculated. The previous step was repeated for all slices along the masked image and the mean of all FWHMs was recorded as the chromosome width. In the final step, the chromosome widths were halved to obtain chromosome arm widths.

At the 30-minute and later time points, chromosome arms were traced by manual shaping in Fiji. Ellipsoid approximation was then applied in Fiji, which generates an ellipsoid of the same area as the traced shape. The lengths of the primary and secondary ellipsoid axes were recorded as chromosome arm lengths and widths. Roundness was calculated by dividing the widths by the lengths. Regression lines were plotted using the “stat_smooth” function in ggplot2.

### Simulations of chromatin chain behavior

Simulations were performed using a previously described chromatin simulation package^21^, where briefly, the simulation consists of Brownian dynamics of beads connected by springs, with soft mutual repulsive interactions, such that with no further interactions the chain is a self-avoiding Rouse polymer. Each bead effectively corresponds to a radius of 25 nm, incorporating approximately 10 nucleosomes and roughly 2 kb of DNA. Therefore, simulations in this report, between *N* = 100 and *N* = 4000 beads, correspond to chromatin chains roughly 200 kb to 8 Mb in length. Without added loop capture interactions, theses simulations are in effect “random walk” simulations, and they are initialized using a random walk configuration with random bond angles between beads on a unit sphere.

In addition, we can turn on “loop-capture” interactions during these simulations, mediated by condensin binders that are found every 10 beads, roughly corresponding to the average frequency of condensin binding sites in fission yeast^16,21^. A small change was made to the simulation code, such as to allow random elongated ellipsoidal initial conditions, which were used for the loop capture simulations. Here, beads are placed randomly along the *z*-axis by drawing from a Gaussian distribution of standard deviation *L*_0_/2, where *L*_0_ is the specified initial length of the polymer. The *z*-axis positions are then sorted by position, after which each of these beads is assigned a random transverse *x* and *y* position, by drawing from a Gaussian distribution of standard deviation *w*_0_/2, where *w*_0_ is the specified initial width and depth of the polymer; since our data cannot distinguish width and depth, we let them be initially equal in our simulations. The values of *L*_0_and *w*_0_we choose for each value of *N* the number of beads in the simulation – are roughly 4 x the equilibrium lengths and 1 x the equilibrium width, which we found from test simulations for each bead length. For simulations testing a different initial condition (Supplementary Fig. 4), we used 1 x the equilibrium lengths and 1 x the equilibrium width, for *L*_0_ and *w*_0_, respectively.

The length *L*, width *w*, and depth *d*, for any conformation of the polymer in simulation is determined by calculating the eigenvalues of the covariance matrix of bead positions; if *λ*_1_, *λ*_2_, *λ*_3_ are these eigenvalues ordered by decreasing magnitude (*λ*_1_ > *λ*_2_ > *λ*_3_) then 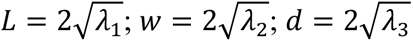.

Exponential fits on simulated data were performed in MATLAB, by first using “fminsearch” – which uses the Nelder-Mead simplex method – and then using the output fit parameters as the initial condition for non-linear regression using the “lsqcurvefit” function, which produces standard error estimates on the optimal parameters. Fits are performed on the average length and width from 10 replicate simulations for each polymer length *N* reported. The equilibrium length is one resultant fitting parameter, and it is this value that is reported as the average equilibrium length or width.

### Statistical analysis

To comprehensively measure chromosomes, the dimensions of all chromosomes or chromosome arms from two independent chromosome spreads were recorded at each time. When chromosome width was measured using Gaussian fitting, individual measurements at each pixel offset were aggregated and the mean was recorded as the chromosome width, then halved to approximate chromosome arm width. Power law exponents were derived from the length to width distributions, plotted on double logarithmic scale, by fitting linear regression lines using the “stat_smooth” function in ggplot2. 10 repeat simulations were conducted in all cases. Means and standard deviations of all measurements on simulated chains are reported. Equilibrium dimensions were fitted using non-linear regression using the “lsqcurvefit” function in MATLAB.

## Data, Materials, and Software Availability

All results obtained are included in the article and/or supporting information. The modified MATLAB code used for semiautomatic chromosome width measurements is available at https://figshare.com/s/63ff965dd434286fab21. The code for the biophysical simulation of chromatin chain behavior was described previously^21^ and is available from the GitHub repository (https://github.com/FrancisCrickInstitute/Chromosome-Condensation).

## Acknowledgements

We would like to thank Paul Bates and Eric Kramer for their input, as well as M. Molodtsov and our laboratory members for discussions and critical reading of the manuscript. This work was supported by the Dr. Yoshifumi Jigami Memorial Fund, The Society of Yeast Scientists, the Institute for Fermentation, Osaka (IFO) and Waseda University grants for Special Research Projects (2021C-387, 2022C-306, 2023C-283, 2024C-285, to Y.Ka.), JSPS research grants (22K06092 to Y.Ka., as well as 22H04996 and 24H02283 to T.H.), a Wellcome Trust Investigator Award (220244/Z/20/Z to F.U.), and the Francis Crick Institute that receives its core funding from Cancer Research UK, the UK Medical Research Council, and the Wellcome Trust (cc2137).

## Author Contributions

Y.Ka., B.S.K. and F.U. conceived the study, Y.Ku. and T.H. performed cell synchronization and microscopy, Y.Ka. performed measurements and data analyses, T.C., M.L. and B.S.K. performed the computational simulations and their analyses, B.S.K. and F.U. wrote the paper with input from all coauthors.

---

**Supplementary Figure 1.**
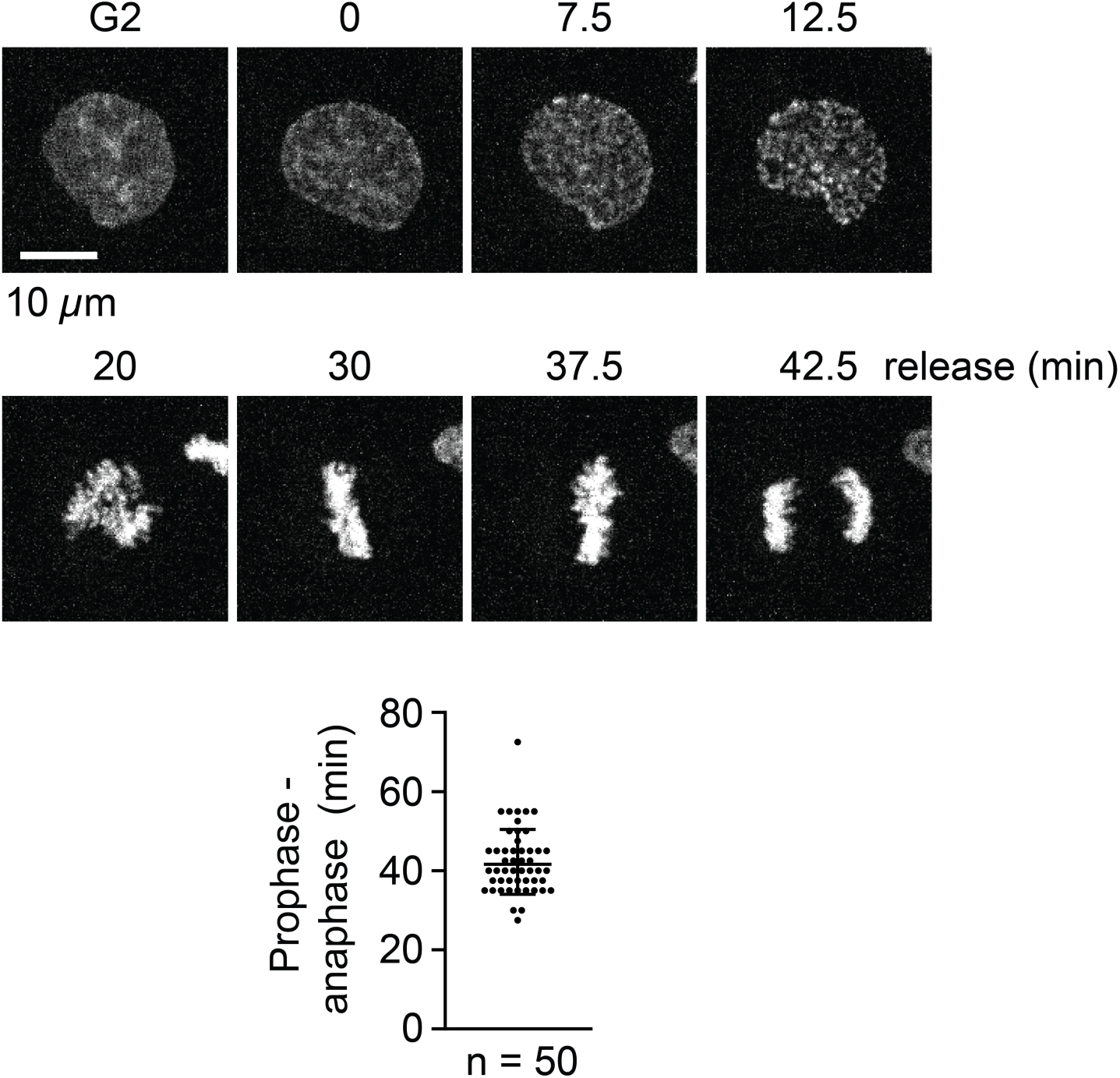
Timing of chromosome segregation without colcemid. Cells were synchronized in G2 and released, as in the experiment shown in Figure 1, but in the absence of colcemid. A time series of images of a cell traversing though mitosis is shown. The time of anaphase onset was determined in 50 cells. Each measurement is shown, together with the mean and standard deviation.

**Supplementary Figure 2.**
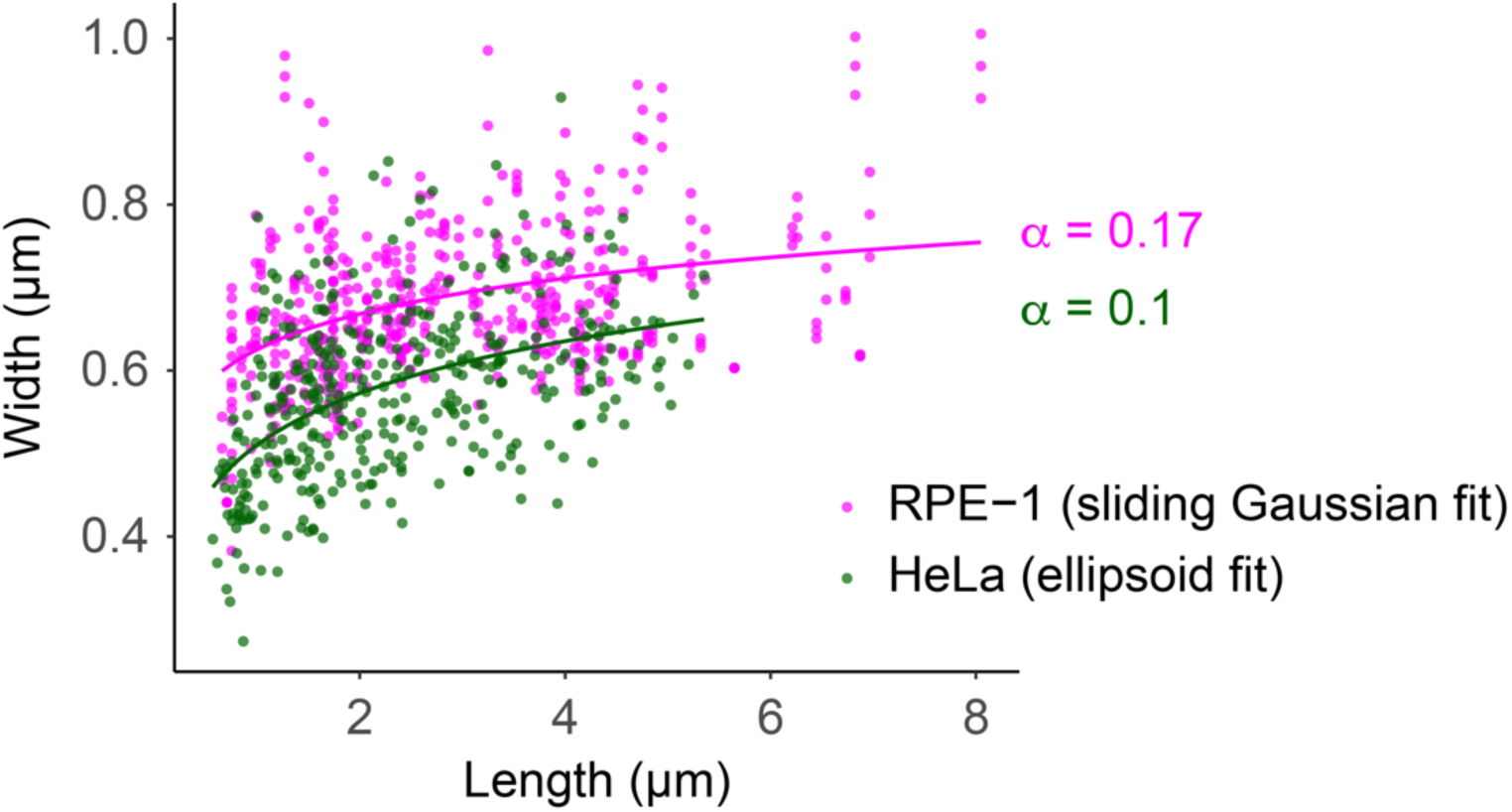
Comparison of chromosome width measurements by ellipsoid fitting with those obtained using a sliding Gaussian fitting algorithm. Chromosome width measurements obtained 30 minutes after release of human RPE-1 cells from a G2 arrest using a sliding Gaussian fitting approach, which traces straight sections of chromosome arms (pink) ^11^, are compared with measurements obtained in our current study using similarly synchronized HeLa cells and an ellipsoid fitting approach (green). We use different cell lines and different measurement methods and, as expected, ellipsoid fitting appears to have underestimated some chromosome lengths. While both sets of measurements are therefore quantitatively different, they both reveal that, qualitatively, longer chromosome arms are wider. This arm length to width relationship can in both cases be fitted by power laws with comparable exponents α.

**Supplementary Figure 3.**
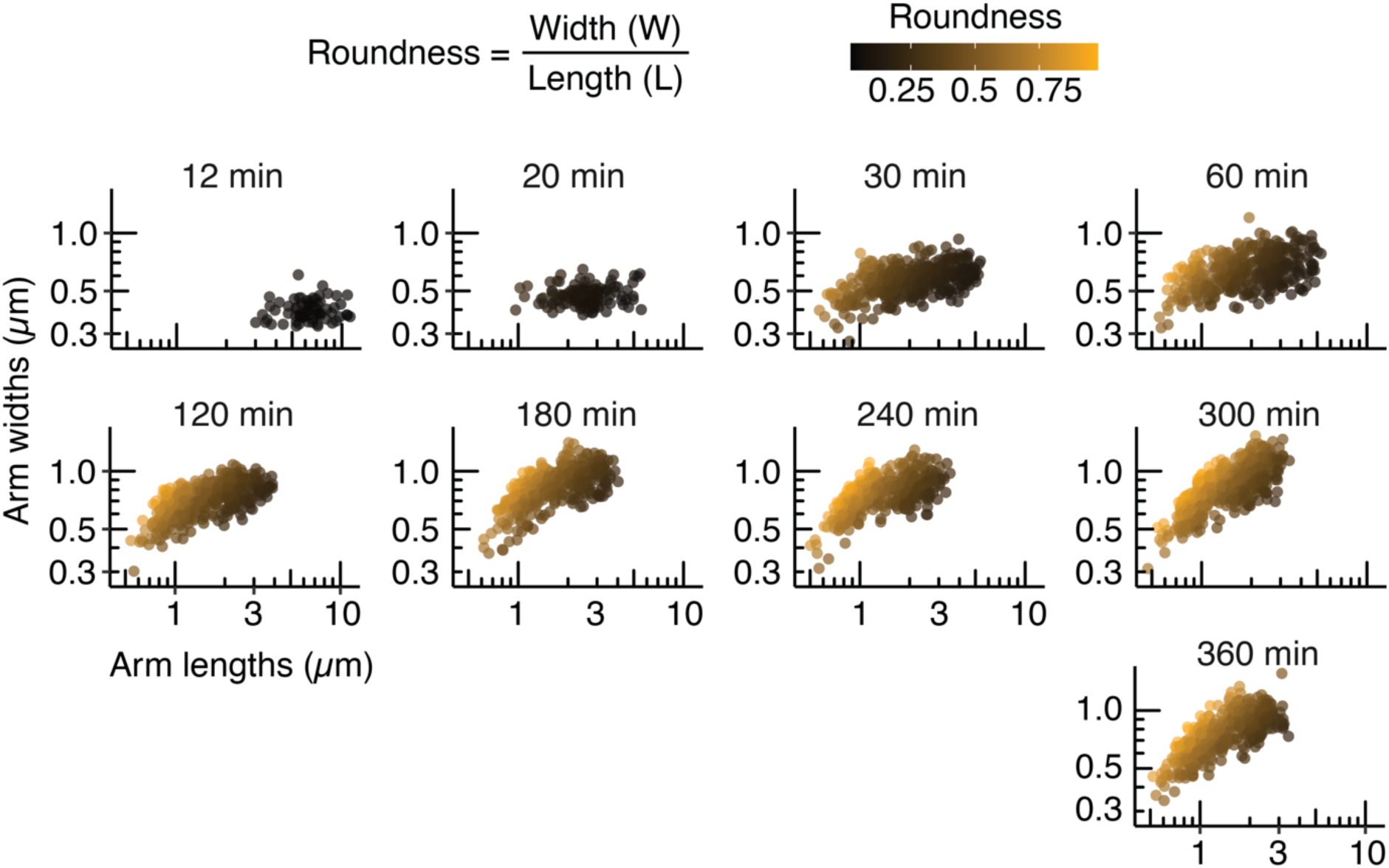
Chromosome roundness over time. Chromosome width as a function of length, at the designated times are shown, with increasing roundness indicated as hues of black to yellow.

**Supplementary Figure 4.**
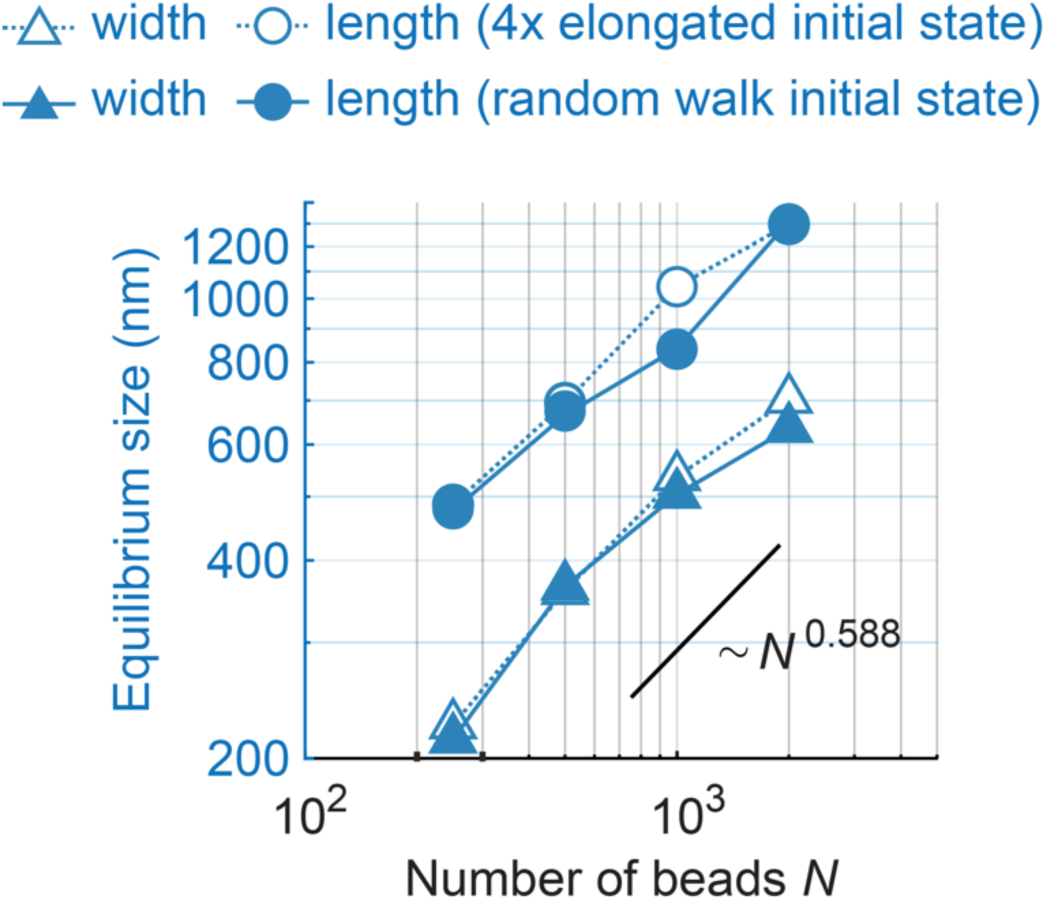
Comparison of the impact of two different initial states on the equilibrium state of loop capture polymer simulations. Equilibrium chromosome lengths and widths, when simulations of increasing chain lengths were started from either an unstretched random conformation, or from a 4 x elongated conformation are shown. The theoretically expected scaling behavior of random self-avoiding polymers is indicated for comparison.

